# Glia-neuron signaling mediated by two different BMP ligands impacts synaptic growth

**DOI:** 10.1101/2021.02.23.432606

**Authors:** Mathieu Bartoletti, Tracy Knight, Aaron Held, Laura M. Rand, Kristi A. Wharton

## Abstract

The nervous system is a complex network of cells whose interactions provide circuitry necessary for an organism to perceive and move through its environment. Revealing the molecular basis of how neurons and non-neuronal glia communicate is essential for understanding neural development, behavior, and abnormalities of the nervous system. BMP signaling in motor neurons, activated in part by retrograde signals from muscle expressed Gbb (BMP5/6/7) has been implicated in synaptic growth, function and plasticity in *Drosophila melanogaster*. Through loss-of-function studies, we establish Gbb as a critical mediator of glia to neuron signaling important for proper synaptic growth. Furthermore, the BMP2/4 ortholog, Dpp, expressed in a subset of motor neurons, acts by autocrine signaling to also facilitate neuromuscular junction (NMJ) growth at specific muscle innervation sites. In addition to signaling from glia to motor neurons, autocrine Gbb induces signaling in larval VNC glia which strongly express the BMP type II receptor, Wit. In addition to Dpp’s autocrine motor neuron signaling, Dpp also engages in paracrine signaling to adjacent glia but not to neighboring motor neurons. In one type of dorsal midline motor neuron, RP2, *dpp* transcription is under tight regulation, as its expression is under autoregulatory control in RP2 but not aCC neurons. Taken together our findings indicate that bi-directional BMP signaling, mediated by two different ligands, facilitates communication between glia and neurons. Gbb, prominently expressed in glia, and Dpp acting from a discrete set of neurons induce active Smad-dependent BMP signaling to influence bouton number during neuromuscular junction growth.

## INTRODUCTION

The efferent or motor nervous system enables signals from the CNS to invoke an action in peripheral organs, such as the contraction of muscles resulting in locomotion or other motor response. Intimately tied to an efferent response is the afferent or sensory response that brings sensation back to the central nervous system. The simple motor circuit incorporates elements of each, motor neurons, sensory neurons, and the target tissue, muscle fibers. In addition, non-neuronal glial cells are critical for the proper formation and function of this neural circuit. They form a protective blood brain barrier, and while glia do not themselves produce electrical impulses, they have a profound impact on neurotransmission by modulating synaptic development, maintenance and efficacy, as well as by providing metabolic support to neurons and instructional cues for axon guidance (Auld and Robitaille 2003; Kanai et al. 2018; Volkenhoff et al. 2015).

Inherent in the formation of neural circuits is the highly-controlled communication between the various cellular components, i.e., neuron to neuron, neuron to muscle, muscle to neuron, glial cell to neuron. The directionality of intercellular signaling depends on the source of a secreted ligand or signal, and the localization of receptors in the membrane of receiving cell(s). In response to ligand-receptor interactions extracellular space, intracellular effectors transduce the signal by altering the receiving cell’s physiology. The BMP/TGF-β (bone morphogenetic protein/transforming growth factor-β) signaling pathway plays critical roles in facilitating communication between different components of the motor circuit in both *Drosophila* and vertebrates (Bakkers et al. 2002; Dudas et al. 2004; Kokubu et al. 2018; Matsuura et al. 2008; Tsujii et al. 2009; Zhang et al. 2002). Ligands of the BMP/TGF-β superfamily are processed from homo- or heterodimeric proproteins as it transits the golgi, and secreted from the source cell. Active ligands bind heterotetrameric receptor complexes composed of type I and type II transmembrane serine/threonine kinase receptors. In canonical BMP signaling, the constitutively active type II kinase phosphorylates to activate the type I receptor upon ligand binding to the receptor complex. In response, the receptor-mediated Smad cytoplasmic signal transducer, R-Smad, is phosphorylated enabling its accumulation in the nucleus where it elicits a transcriptional response. In the case of canonical BMP signaling in Drosophila, nuclear accumulation of the phosphorylated Smad1/5/8, pMad, is a useful indicator of active signaling in that cell (reviewed in (Raftery and Sutherland 1999)). Autocrine, as well as paracrine BMP signaling occurs, the distinction of which depends on the specific removal of the ligand from the source cell, and observation of signaling output in the receiving cell. Identifying the source of a signal and its consequences on target cells is not always trivial. Loss of function analyses provide important information about function, and given the non-autonomous nature of most secreted ligands such as BMPs, it is especially important to recognize that over- or misexpression of ligand genes highlight where signal reception is possible, but does not necessarily reflect signaling capabilities of the endogenous source. Similarly, in overexpression studies, the inherent specificity of BMP ligands is lost as different ligands are no longer subject to normal regulatory mechanisms.

Multiple signaling components have been implicated in neural development in vertebrates as well as in Drosophila. In Drosophila, a number of studies to date have characterized roles for BMP signaling in the development of the nervous system, as well as in ensuring its proper function. *glass bottom boat* (*gbb*), the gene encoding the *Drosophila* BMP5/6/7/8 orthologous ligand, Gbb (Wharton et al. 1991), has a demonstrated role in both growth and function of the larval neuromuscular junction (NMJ), as well as synaptic plasticity (Berke et al. 2019; Berke et al. 2013; Goold and Davis 2007; Hoover et al. 2019; James and Broihier 2011; James et al. 2014; McCabe et al. 2003; Sulkowski et al. 2014). Active BMP signaling is first evident in the embryonic nervous system at stage 15 with the nuclear localization in a subset of neurons of pMad, the phosphorylated Drosophila R-Smad1/5/8, an indication that these neurons have received a BMP signal (Marques et al. 2002). pMad is lost from the nervous system in embryos null for *gbb*, or for *wishful thinking* (*wit*), the gene encoding the BMPR2 type II receptor ortholog (Marques et al. 2002; McCabe et al. 2003). *gbb* is primarily expressed in the mesoderm and invaginating midgut primordia of the embryo (Doctor et al. 1992), while *wit* is expressed in a specific subset of neurons in the CNS during embryogenesis (Aberle et al. 2002; Marques et al. 2002). The loss of pMad in *wit* mutant but not in mutant embryos for punt, another type II receptor gene, indicates a specific requirement for Wit, in addition to the Gbb ligand. A role for type I receptors Tkv and Sax has been suggested in the embryonic CNS (Marques et al. 2002; McCabe et al. 2003), and the reduced number of boutons in NMJ6/7 of *sax* mutant larvae, with a reduction in the amplitude of excitatory junction potentials (EJP) and a lower mean quantal content demonstrates a role for the Sax receptor in larval NMJ growth and function (Rawson et al. 2003).

*gbb* and *wit* are also required for normal NMJ structure and neurotransmission in the larva (Aberle et al. 2002; Lee et al. 2016; Marques et al. 2002; McCabe et al. 2003; Piccioli and Littleton 2014). While pMad staining remains in the nuclei of distinct sets of neurons in the larval ventral nerve cord (VNC), neuronal specific staining of Wit is lost and its expression becomes diffuse during the larval stage (Aberle et al. 2002; Marques et al. 2002). The defects in synaptic growth and function evident in *gbb* mutant larvae have been attributed to two somewhat separable functions. Targeted overexpression of *gbb* in the muscles of *gbb* null animals rescues NMJ size, implicating Gbb as a retrograde signal from muscle to motor neuron as a promoter of synaptic growth. Consistent with this role, the knock down of *gbb* in muscles of wild type animals results in small NMJs similar to that seen in wit mutants (Aberle et al. 2002; Berke et al. 2013; James and Broihier 2011; McCabe et al. 2003). A retrograde Gbb signal has also been shown to activate motor neuron expression of *trio,* a Rac and Rho guanine nucleotide exchange factor with a role in regulating synaptic growth through a pathway controlling actin dynamics (Ball et al. 2010; Pawson et al. 2008). On the other hand, targeted overexpression of *gbb* in neurons is less effective at increasing NMJ size, but is able to restore compromised synaptic transmission (Berke et al. 2013; Goold and Davis 2007; James and Broihier 2011; McCabe et al. 2003). A proposed role for a neuronal pool of Gbb in neurotransmitter release emerged most recently from experiments in which expressing *gbb* in neurons results in trafficking of Gbb in dense core vesicles with neuronal factor Crimpy (Cmpy) (James et al. 2014) and the neuronal expression of *gbb*-RNAi in a *gbb* null heterozygote phenocopies the synaptic defects. associated with calcium channel α_2_δ-3 mutants (Hoover et al. 2019). In its earlier requirement for synapse growth and later as a facilitator of neurotransmission, the activation of BMP signaling by Gbb involves pMad-mediated signaling (Berke et al. 2013; Hoover et al. 2019).

Here, we report that two BMP ligands contribute to synaptic growth from different sources, and both engage in glia-neuron signaling. As in vertebrates, a large portion of the non-neuronal component of the motor system in *Drosophila* is composed of a variety of different types of glia, each with a distinct function, conserved with their vertebrate counterparts (Bittern et al. 2020; Freeman and Doherty 2006; Ou et al. 2014; Stork et al. 2012). We find that *gbb* is expressed in several types of glia, as well as the body wall muscles and VNC associated trachea of third instar larvae, in addition to its previously established expression in embryonic mesoderm, the larval dorsal neurohemal organs and neuroblasts located in the larval brain lobes (Doctor et al. 1992; Kanai et al. 2018; Marques et al. 2003; McCabe et al. 2003; Staehling-Hampton et al. 1994). Glial-expressed *gbb* is required for normal synaptic growth at the NMJ, highlighting glia as a source of Gbb signals important in motor neuron development. Reduced NMJ growth associated with knock down of *gbb* in glia is accompanied by a strong reduction in pMad in motor neuron nuclei, indicating that Smad-mediated signaling from glial cells to motor neurons plays a role in proper synapse development. The Drosophila BMP2/4 ortholog, Dpp, encoded by *decapentaplegic* (*dpp*), is not detected in glial cells, but rather in a subset of motor neurons whose cell bodies reside in the dorsal and dorsal lateral VNC. Knocking down *dpp* in these neurons leads to a cell autonomous reduction in pMad with an associated reduction in the synapses of these motor neurons at their target muscles. Furthermore, Dpp-dependent autocrine signaling preferentially regulates *dpp* expression in RP2 motor neurons via a positive feedback loop. These findings highlight the role of BMPs from distinct sources in glia-neuron communication and underscores their respective importance in synaptic growth.

## MATERIALS and METHODS

### Fly Stocks and culture

Flies were raised at 25°C on a standard cornmeal/sugar/agar food. For all experiments 3 virgin females were crossed to 2 males and brooded daily to control for development. Both sexes of the experimental and control classes were used for the experiments. *w; gbb-Gal4/Cyo #1* and *w; gbb-Gal4#5* were derived from the *gbbR-gbb* lines described in Kanai et al 2018. *w; UAS-mcd8GFP/CyO, w; Repo-Gal4/TM6c* (BSC#7415)*, w; GFP-RNAi, w; UAS-gbb-RNAi, UAS-dicer2 (*abbreviated in the text in UAS-*gbb-RNAi)* (Ballard et al. 2010)*; UAS-gbb* (9.7 line, Khalsa et al. 1998)*; w; dpp-blk-Gal4, UAS-GFP/T2-3, w; shdpp2HB* (gift from Michael Levine, abbreviated in the text in UAS-*dpp-RNAi*), Wit-2xTY1-sGFP-V5-BLRP-3x-FLAG (VDRC#318043) (Sarov et al. 2016) were the other lines used.

### Dissections and immunofluorescence

Wandering third larval instars were selected as larvae >108 hours after egg lay. The larvae crawled out of the food and had not everted their anterior spiracles. Larvae were washed and then dissected in fresh PBS 1X (Sigma P4417). During dissection, each larva was pinned on an agarose plate in the head and in the tail. The dorsal side was cut alongside the anterior to posterior axis. Both side of the cut were opened and pinned to the plate before the interior of the larva was cleaned from all content non-related to the nervous system, leaving the VNC, the brain and the muscle carrying the NMJ accessible for immunochemistry and microscopy. Each larva was fixed in PBS1X + 4% paraformaldehyde (16% paraformaldehyde Electron Microscopy Sciences 15710) for 20 min at room temperature. Larvae were next washed 3 times in PBS1X + 0.3% triton (Sigma T8787) for 10 min at room temperature. Larvae were then blocked in PBS1X + 0.3% triton + 1% NGS (Jackson Immuno Research 005-000-121) for 1 hour at room temperature and incubated over-night at 4°C in a solution of primary antibodies diluted in the same solution as the one used for blocking. Samples were next washed 3 times in PBS1X + 0.3% triton for 10 min at room temperature and incubated in a secondary antibody solution for 2 hours at room temperature or over-night at 4°C. As for the primary antibodies solution, secondary antibodies are diluted in the blocking solution. After 3 new washes in PBS1X + 0.3% triton, samples were incubated for 10 min at room temperature in a PBS1X + 0.3% triton + Hoechst (Molecular Probe 5ug/ml). Samples were then mounted in 80% glycerol + 0.5% N-propyl-gallate (Sigma-Aldrich) and imaged with a ZEISS LSM500 or ZEISS LSM800 or Olympus FV3000 confocal microscopes. For a better homogeneity during the immunostaining treatment, the larvae from the different genotypes were pulled in a same Eppendorf tube. Depending on their genotype, each larva was left intact or had a cut in its most posterior part, allowing to recognize the genotype of each dissection. No more than 10 dissected larvae (test + control) were pooled together in the same Eppendorf to make sure that each sample could be in contact with the different reagent necessary for immunostaining. Each experiment was repeated independently at least two to three times. Primary antibodies and stains used in this study are: mouse anti-Dlg (DSHB, 4F3, RRID:AB_528203 at 1/200), rabbit anti-HRP-647 (Jackson Immuno Research, RRID:AB_2338967 at 1/100), rabbit anti-pSmad3 (Abcam ab52903, RRID:AB_882596 at 1/500), mouse anti-GFP (REF at 1/200), rat anti-ElaV (DSHB, 7E8A10, RRID:AB_528218 at 1/100), mouse anti-Repo (DSHB, 8D12, RRID:AB_528448 at 1/50), mouse anti-Flag (Sigma F3165), phalloidin-568 or phalloidin-488 (Life Technology A12380 and A12379 at 1/100). The secondary antibodies used in this study are: goat anti-rabbit 568 (Life Technology A11011 at 1/200), goat anti-rat 647 (Life Technology A21247 at 1/200), goat anti-mouse 488 (Life Technology A11001 at 1/200), goat anti-mouse Cy3 (Life Technology A10521 at 1/200), goat anti-chicken 488 (Life Technology A11039 at 1/200) and goat anti-rabbit 488 (Cell Signaling Technology #4412 1/200).

### Electrophysiology

Wandering 3rd instar larvae were dissected in modified HL3.1 containing (in mM): 70 NaCl, 5 KCl, 10MgCl_2_, 10 NaHCO_3_, 5 trehalose, 115 sucrose, 5 HEPES, 0.5 CaCl_2_ and pH 7.2. Larvae were fileted to expose the body wall and the VNC and organs were removed. Sharp electrodes (20-25MΩ) were filled with 3M KCl and used to record from muscle 6 in segment A3. Only muscles with an input resistance >5MΩ and a resting membrane potential of <-60mV were used. A 1nA current was applied for 500ms for 50 sweeps to determine the muscle resistance and capacitance values. mEPSPs were recorded for 3 minutes in the absence of nerve stimulation. eEPSPs were evoked by stimulating axons innervating muscle 6 with a 0.3ms square pulse of current from a suction electrode. A minimum of 10 eEPSPs were evoked and then averaged to determine the eEPSP value for that animal. Recordings were made using an Axopatch 200B, sampled at 10khz, filtered at 3khz, digitized with a Digidata 1440A, and analyzed using custom MATLAB scripts (Held et al. 2019). To compare 2 genotypes, we used a student’s t-test accounting for the variance.

### Bouton counting

Boutons were visualized with a Zeiss AXIO imager.M1 compound microscope using a 63x objective. Boutons were counted by using the anti-Dlg staining on NMJ innervating muscle 6/7, 4, 3 or 2 on both A3 hemisegment. We averaged the number of synaptic boutons that were counted on each hemisegment, and come from a similar type of muscle. That average is used as the bouton number value for each larva. To compare 2 genotypes, we used a student’s t-test accounting for the variance.

### pMad quantification

To compare potential differences between p-Mad fluorescence across different genotypes we kept the same confocal settings when control and experimental VNC pictures were acquired.

#### Motor neurons from the midline cluster

The p-Mad fluorescence intensity present in the motor neurons nuclei located at the VNC midline at segment A6 was quantified using the measure tool from ImageJ. An anti-ElaV nuclear-specific antibody was used to delimit the surface occupied by each neurons nucleus and then applied to the p-Mad channel allowing the exact quantification of p-Mad fluorescence. The value of 10 to 12 neurons was quantified from each sample were then pooled according to their genotypes. Their average was compared using a student’s t-test accounting for the variance.

#### Peripheric neurons

As for the midline cluster, we made use of the anti-elaV staining to select all the neuron nuclei present on the periphery of the midline cluster. The surface occupied by all the peripheric neurons was then applied to the p-Mad channel to quantify p-Mad fluorescence. Each value was pooled according to genotype. Their average was compared using a student’s t-test accounting for the variance.

#### Glial cells

The p-Mad fluorescence intensity present in the glial nuclei located near the VNC midline at segment A6 was quantified using the measure tool from imageJ. An anti-elav nuclei specific antibody was used to counter select Glial nuclei and Hoechst was used to delimit the surface occupied by each glial nucleus and then applied to the p-Mad channel allowing the exact quantification of p-Mad fluorescence. The value of 6 glial cells was quantified from each sample and then pooled according to their genotypes. Their average was compared using a student’s t-test accounting for the variance.

#### RP2, aCC and U GFP-positive neurons and other non GFP neurons

The same method using the anti-ElaV staining was performed to quantify p-Mad fluorescence in these neurons. The values where then separated according to their location and/or their ability to express GFP under the control of the *dpp-blk>* driver and pooled in a RP2 and aCC motor-neuron (GFP positive MN located in the midline of the A5 segment), U neurons (GFP positive set of neurons located on the side of the midline at the A2 segment) and *dpp* negative motor neurons (GFP negative neurons located in the midline of the A5 segment). Averages were compared using a student’s t-test accounting for the variance.

### *Dpp-blk>GFP* quantification

GFP fluorescence levels was quantified in both aCC and both RP2 of segment A5 with ImageJ. For each image, both value for aCC or RP2 were then averaged to give a single value representing the GFP fluorescence in each neuron type. Because the confocal channel used to acquire GFP fluorescence signal was not using the same settings between the different pictures taken we could not directly compare GFP fluorescence values between genotypes. To counter this issue, we did an aCC/RP2 GFP fluorescence ratio for each picture. These ratios were averaged for each genotype and were compared using a student’s t-test accounting for the variance.

## Data availability statement

The authors affirm that all the data needed to support the conclusions presented in this article are present in the figures and the text of the article. Drosophila strains are available upon request.

## RESULTS

### *gbb* is expressed in multiple types of glia in the VNC

As part of our ongoing studies on *gbb* function, we sought to better understand the sources of Gbb signals with regard to the larval motor circuit. We generated a *gbb-Gal4* transgene by replacing the *gbb* coding sequences contained within a *gbbR-gbb* genomic rescue construct with the yeast transcriptional activator GAL4 (Kanai et al. 2018)(Figure 1A). Independent insertions, mediated by P-element-mediated transposition of the *gbb-Gal4* transgene, at random sites in the *Drosophila* genome were recovered. This construct, with GAL4 under the control of genomic *gbb* cis-regulatory sequences, enables assessment of the endogenous pattern of *gbb* expression when crossed to *UAS-mcd8-GFP*, a membrane bound GFP (Figure 1, S1). In fileted preparations of *gbb-Gal4;UAS-mcd8-GFP* wandering third instar larvae, the *gbb>mcd8-GFP* signal is evident as expected in body wall muscles, as well as in the neurohemal organs located along the dorsal midline of the ventral nerve cord (VNC) (Figure 1B,E; Supplementary Figure 1B,D; Marques 2003). We found *gbb>mcd8-GFP* signal clearly visible in several types of glial cells, those in the segmental nerves that wrap the axonal projections of motor and sensory neurons, and a subpopulation of glia that encase the VNC (Figure 1 B-D,F). When compared with the pattern of GFP in all glial cells (repo>mcd8-GFP (Figure S1A), those expressing *gbb>mcd8-GFP* are likely a subset of glia that envelope the VNC (Figure S1), evident in cross sectional images of the VNC (Figure 1C, Figure S1A’,B’). Optical cross sections show that gbb is expressed in wrapping glia that insulate individual axonal projections within the segmental nerve and likely subperineural glia that also encase the VNC (Figure 1F). In addition to glia of the segmental nerves, *gbb* is expressed in a discrete set of thread-like structures with a luminal characteristic of trachea whose nuclei do not express Repo (Figure 1F).

**Figure 1:**
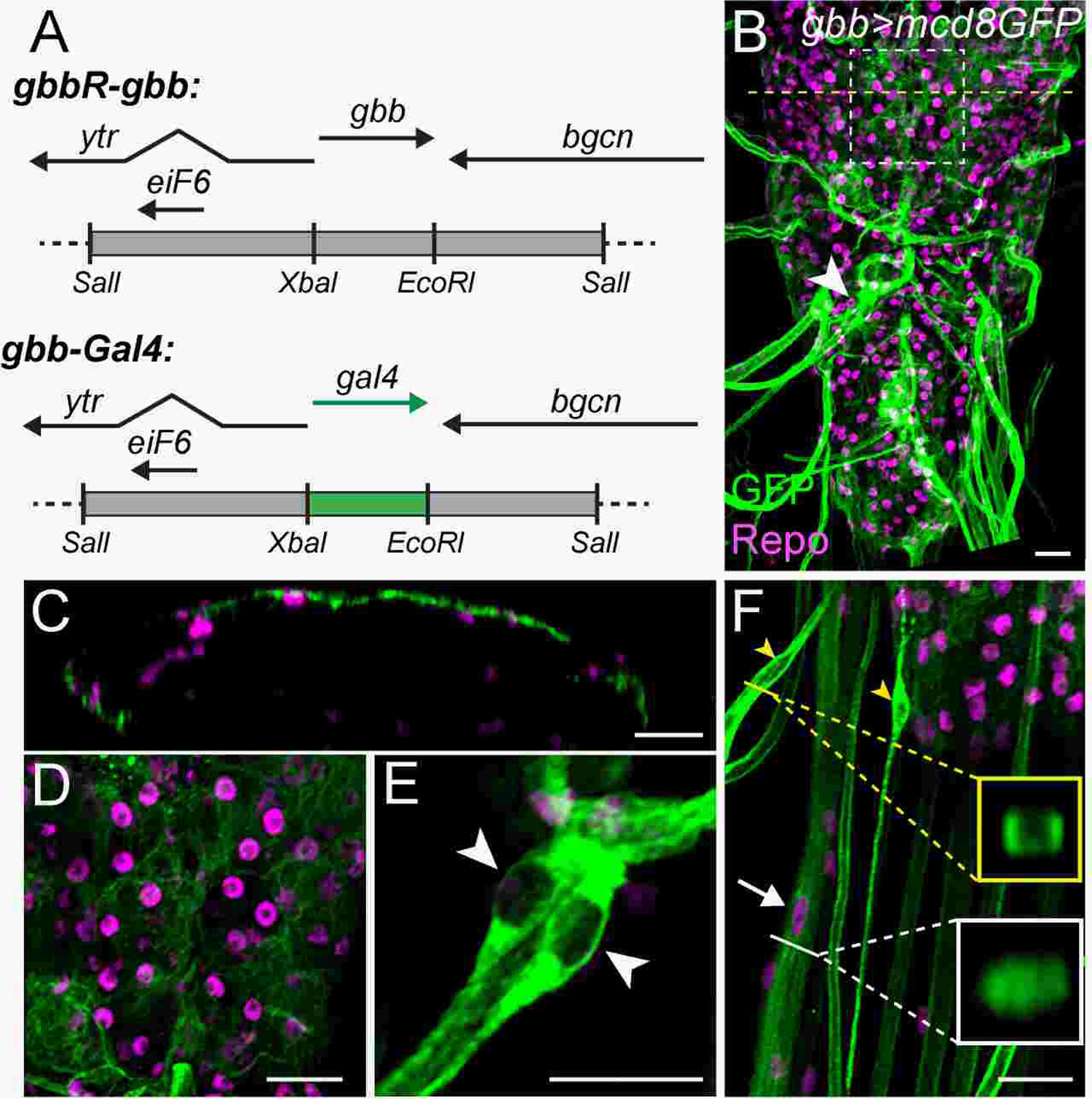
*gbb* is expressed in multiple cell types, including glia, in the *Drosophila* third larval instar ventral nerve cord (VNC). (A) *gbb* coding sequences in the genomic *gbb* rescue construct (gbbR-gbb) were replaced with Gal4 to generate *gbb-Gal4* transgenic lines (Kanai et al 2018). (B-D) Immunohistochemistry of membrane-bound GFP and Repo expression in the ventral nerve cord (VNC) of *gbb-Gal4;UAS-mcd8-GFP* third instar larvae. (B) *gbb* is expressed in multiple cell types in the VNC, including (C,D) glial cells enveloping the VNC also visualized when membrane bound GFP is expressed specifically in glia (Figure S1A,*repo-Gal4;UAS-mcd8-GFP*), (E) the dorsal neurohemal organs, (F) wrapping glia of the segmental nerves and trachea (Figure S1C). Dashed line in (B) shows area of cross-section shown in (C). Dashed box in (B) shows enlarged area in (D). (E) High magnification of two cells forming dorsal neurohemal organs marked by large white arrowhead (B). (F) Posterior tip of VNC, projecting segmental nerves, and trachea from *gbb-Gal4;UAS-mcd8GFP* third instar larva highlight *gbb* expression in wrapping glia (arrow, Repo-positive nuclei), tracheal cells (yellow arrowheads, lack Repo staining). Yellow line and box show site of trachea optical cross section highlighting lumen. White line and box indicate site of segmental nerve optical cross section showing gbb expression in wrapping glia within the bundle of axonal projections. anti-GFP (green), anti-repo (magenta) antibodies. Scale bars = 20um.

### Activation of BMP signaling in motor neurons requires glia-expressed Gbb

BMP signaling is activated in motor neurons along the dorsal midline of the VNC, as well as in peripheral neurons, as indicated by the presence of the phosphorylated-Mad signal transducer (pMad), the Drosophila ortholog of Smad1,5,8 (Marques et al. 2002; McCabe et al. 2003; Raftery and Sutherland 1999; Sulkowski et al. 2016). To test if Gbb produced in glia contributes to BMP signaling activation in the VNC, we use the glial enhancer trap driver *repo-Gal4* (Sarov et al. 2016) to knock down *gbb* in glia by expressing *UAS-gbb-RNAi* (*repo-Gal4*; UAS-*gbb-RNAi*) (Figure 2). In the VNC from third instar larvae with *gbb* knocked down in glia, a significant decrease is observed in the level of pMad in nuclei of midline motor neurons (Figure 2C-D’). Quantification of pMad signal intensity in a cluster of 10 midline motor neurons that innervate the sixth abdominal (A6) segment, shows a 56% reduction in pMad levels in *repo>gbb-RNAi* compared to the *repo>GFP-RNAi* control (Figure 2 A-E, boxed cluster, n=8 VNCs for experimental and control, p=1.14E-08). Neurons whose cell bodies reside in more lateral positions of the VNC also show a decrease in pMad levels in *repo>gbb-RNAi* VNCs (53.5%) (Figure 2 A-E, n=8 VNCs for experimental and control, p=0.03). These data indicate that the expression of *gbb* in glia is required for BMP signaling activation in neurons, supporting a role for the Gbb ligand as a mediator of glia to neuron communication.

**Figure 2:**
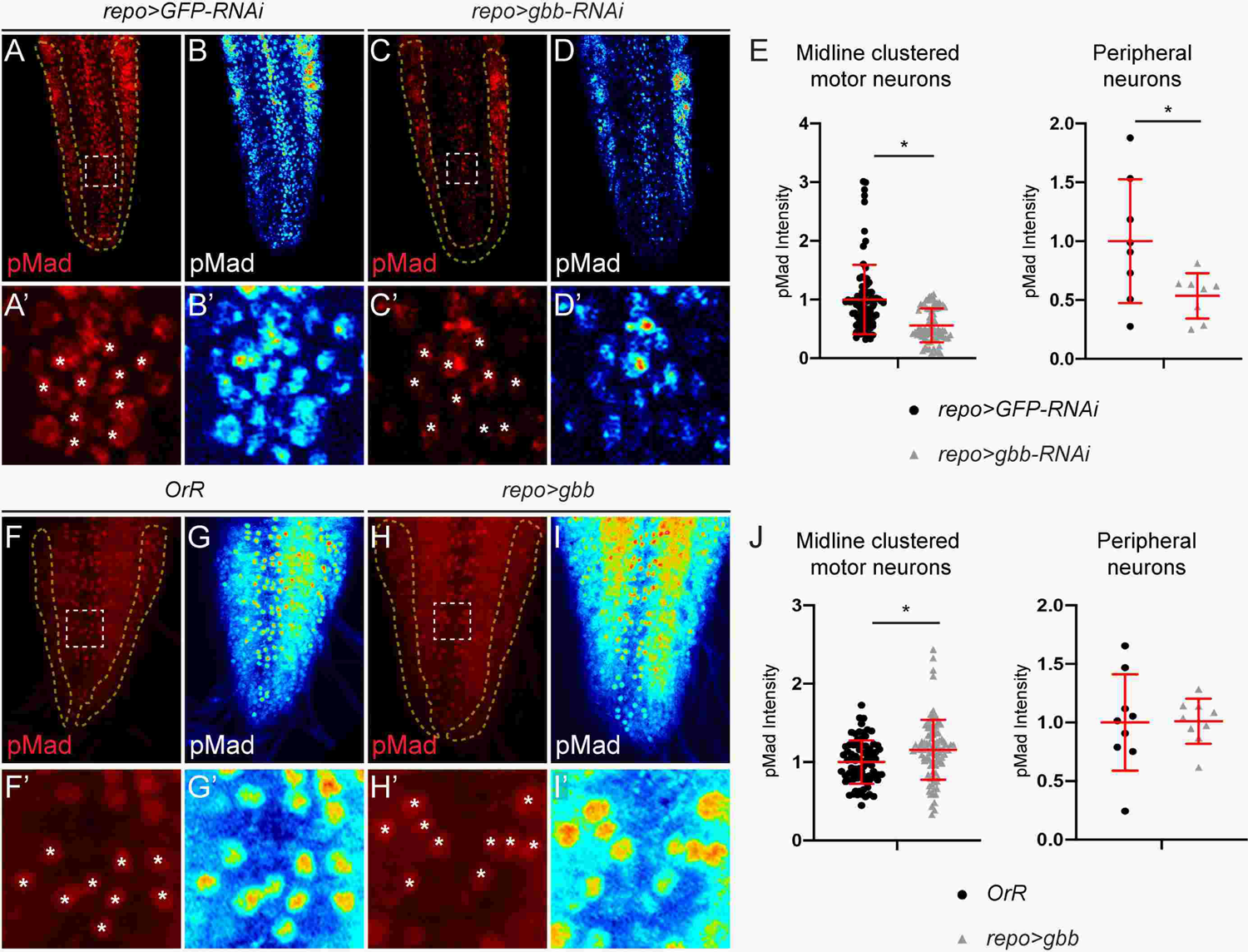
Glial Gbb is required for BMP signaling activation in neurons. (A-D) pMad immunofluorescence signal in VNCs from third instar larvae. (A-B’) control, *repo-Gal4>UAS-GFP-RNAi* and (C-D’) knock-down of *gbb* in glia, *repo-Gal4>UAS-gbb-RNAi,UAS-dicer2*. (A, A’, C, C’) Anti-pMad signal shown in red, or (B, B’, D, D’) in royal color lut representing signal intensity from black (no signal) to red (high intensity). White dashed box in (A) and (C) outline the cluster of midline dorsal motor-neurons innervating segment A6 muscle targets shown at high magnification in (A’, B’, C’, D’). (A’, C’) White asterisks mark location of motor-neurons. pMad signal in peripheral neurons quantified within area outlined by yellow dashed line. (E) pMad fluorescent signal quantification from each of midline neuron within cluster innervating A6 muscles (left) and VNC peripheral neurons (right). Mean intensity and standard deviation (red horizontal lines). T test, * indicates statistical significance at p<0.05. (F-I’) Third larval instar VNC of *OrR* control (F-G’) or overexpression of *gbb* in glia*, repo-Gal4>UAS-gbb* (H-I’). (F, F’, G, G’) Anti-pMad signal in red or (H, H’, I, I’) royal color lut. (J) Quantification of pMad fluorescent signal from each midline neuron within cluster innervating A6 muscles (left), and peripheral VNC neurons (right).. Mean intensity and standard deviation (red horizontal lines). T test, * indictaes statistical significance at p<0.05.

Consistent with the conclusion Gbb can signal from glial cells to neurons, overexpression of *gbb* in glia (*repo-Gal4; UAS-gbb*) results in a 116% increase in pMad levels in the midline motor-neuron cluster (Figure 2F-J, boxed motor neuron cluster innervating A6 muscles, n=8 VNCs for experimental and n=9 for control, p=0.002). No significant change in pMad levels was detected in the lateral regions of the VNC.

### Glial-expressed *gbb* is required for NMJ growth

Given that reduced pMad in motor neuron nuclei has been associated with impaired synapse development, we asked if the glial source of Gbb is required for synapse growth. As a standard morphological measure of neuro-muscular junction (NMJ) development, the number of boutons, sites of neuro-transmitter release, were counted at the NMJs that innervate muscle 2, muscle 3, muscle 4, and muscles 6/7, of segment A3 in wandering third larval instar (Figure 3A,B). In *repo>gbb-RNAi* larvae all four NMJs exhibit a reduction in bouton number compared to control animals (*repo>GFP-RNAi*). An average of 56 boutons in NMJ 6/7 of control animals was reduced to 41 boutons in *repo>gbb-RNAi* (Figure 3A, n=8 for controls, n=7 for experimental, p=5E-04). Reductions in synaptic boutons were also significant at NMJ 4, with a decrease in average number of boutons from 28 in controls to 24 in *repo>gbb-RNAi*, at NMJ3 from 17 in controls to 14 in *repo>gbb-RNAi*, and at NMJ2 from 23 in controls to 16 in *repo>gbb-RNAi* (Figure 3A, n=8 for controls, n=7 for experimental, p=0.04, p=0.01, p=7E-04, respectively). When *gbb* is overexpressed in glial cells (*repo>UAS-gbb*), the average number of boutons at each of these NMJs show a significant increase (Figure 3B). In NMJ 6/7, an increase of bouton number from 52 in control animals to 77 in *repo>gbb*, an average of 20 to 23 boutons at NMJ4, an average of 10 to 12 boutons at NMJ3, and an average of 15 to 18 boutons at NMJ2 (p= 2.8E-04, p=0.03, p=0.02, p=0.03, respectively; Figure 3B). These data indicate that 1) the endogenous expression of *gbb* in glia is required for motor neuron synapse structure, and 2) overexpression of *gbb* in glia can stimulate an increase in NMJ growth. It is important to note that decreasing *gbb* expression in glia impacts the nuclear accumulation of pMad in motor neurons and the growth of NMJs, in the context of endogenous sources of Gbb signal from other cells, such as the muscles.

**Figure 3:**
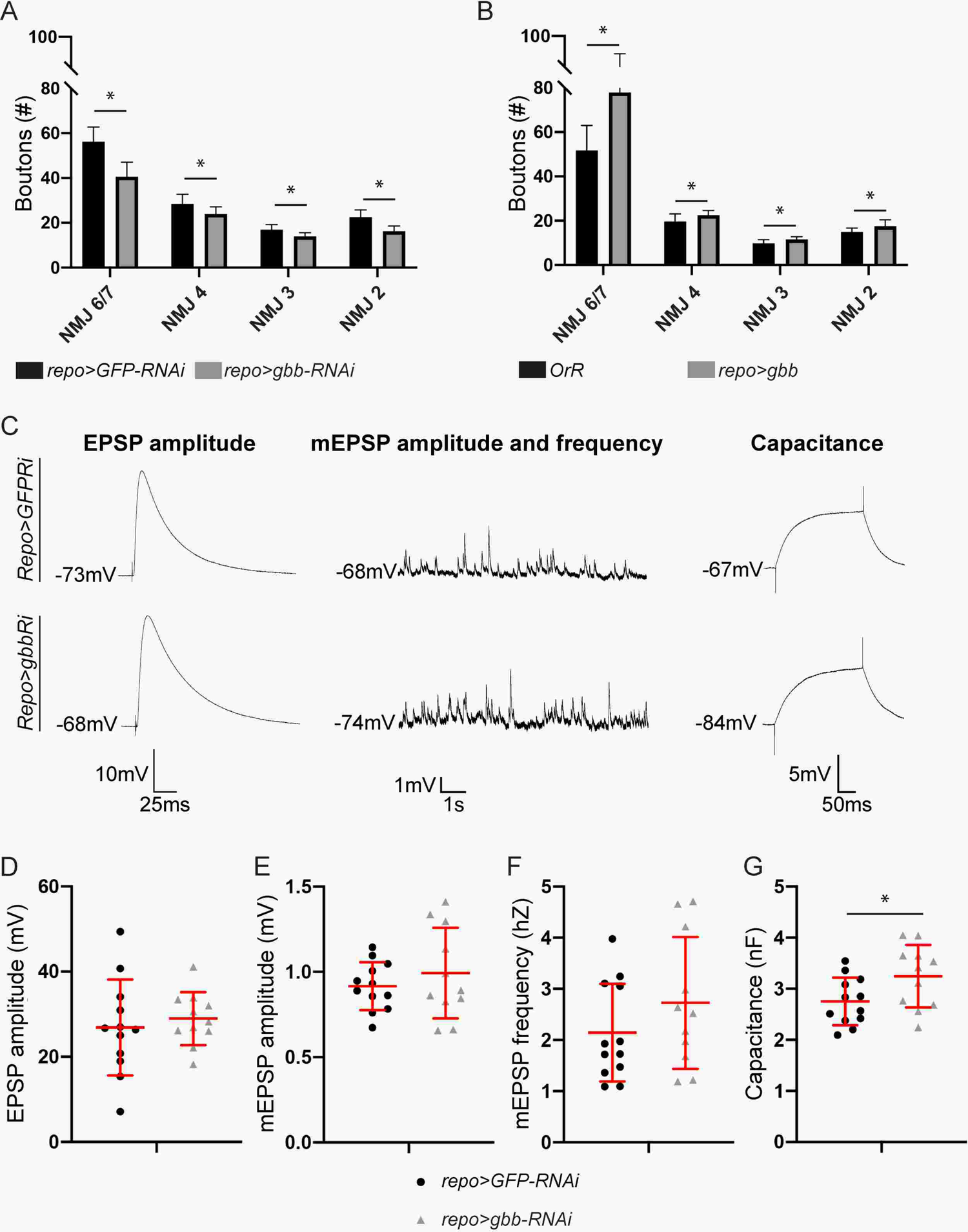
Glial Gbb is required for NMJ growth. (A-B) Bar chart depicting the average number of boutons at NMJ6/7, NMJ4, NMJ3 and NMJ2 in *gbb* knock down *repo>gbb-RNAi, UAS-dicer2* (gray) compared to control animals, *repo>GFP-RNAi* (A) or *gbb* overexpression *repo>gbb* compared to control, *OrR* (B) (black). Error bars denote standard deviation, statistical significance determined by T-test, * indicates p<0.05. (C-F) Electrophysiological recordings from NMJ6/7 at A3 were obtained from *repo>GFP-RNAi* (control) and *repo>gbb-RNAi,UAS-dicer2* wandering third instar larvae. (C) The amplitude of the evoked response (EPSP), as well as the amplitude and release frequency of mEPSPs, and capacitance of the muscle were quantified and showed respectively in (D), (E), (F) and (G). Each data point corresponds to measurement from one larva. Statistical significance determined by T-test, * p<0.05.

We measured the electrophysiological properties of NMJ 6/7 in the A3 segment of *repo>gbb-RNAi* wandering 3^rd^ instar larvae (Figure 3C-G). While modifying *gbb* expression in glia alters the number of boutons (Figure 3A,B), we did not detect a significant change in the amplitude of evoked excitatory postsynaptic responses (EPSP), or of spontaneous miniature excitatory postsynaptic potentials (mEPSP) when *gbb* expression was knocked down in glia (Figure 3C-E). The frequency of minis (mEPSP), while trending higher, did not reach statistical significance in *repo>gbb-RNAi* compared to the control (*repo>GFP-RNAi*) (Figure 3F), but a significant increase in the membrane capacitance of the muscle was evident (Figure 3G average of 3.2nF in *repo>gbb-RNAi* compared to 2.8nF in *repo>GFP-RNAi* control, n=10, p=0.04). This increase in capacitance indicates that the properties of the muscle have been altered as a result of reduced *gbb* expression in glia. Taken together, our findings indicate that *gbb* is expressed in a subset of glial cells, its expression in these cells is required to elicit Smad-dependent BMP signaling in VNC neurons, and it is required for proper NMJ growth and for maintaining the postsynaptic physiology of muscle cells. Importantly, when the expression of *gbb* is down regulated specifically in glia, we detect changes in pMad levels in the cell body of motor neurons, in the bouton number at NMJs, and in the membrane potential of the innervated body wall muscle in the third instar larva, despite the presence of endogenously expressed *gbb* in other cells, sites from which Gbb signals have been implicated in some of these same functions. Down regulation of *gbb* in glia did not affect synaptic properties related to neurotransmission, suggesting that either nuclear pMad in motor neurons is not critical for synaptic function, or that the degree of reduced pMad, is not sufficient to compromise neurotransmission.

### Glial-expressed *gbb* induces autocrine signaling in glia

The ability of Gbb produced by glial cells to invoke signaling in either adjacent or distant cells, is indicative of paracrine signaling. Our analysis of the spatial pattern of BMP signaling activation in the VNC and the segmental nerves revealed that pMad is not only localized to nuclei of neuronal cell bodies but it also accumulates in nuclei of non-neuronal cells of the VNC that do not stain with the pan-neuronal protein ElaV (Figure 4 A,A’), and in peripheral glial cell nuclei identified by anti-Repo staining (Figure S2). To address the possibility that glial-expressed *gbb* may contribute in an autocrine manner to BMP signaling activation in glial cells, we quantified pMad level in glial cells in *repo>gbb-RNAi* compared to controls (*repo>GFP-RNAi*) (Figure 4B-D). The specific knock-down of *gbb* in glia (*repo>gbb-RNAi)* resulted in a 37% decrease in pMad fluorescence intensity in the glial cells nuclei located on either side of the midline motor neuron cluster in the A6 segment compared to *repo>GFP-RNAi* controls (n=6, p=1.11e-05; Figure 4D). This decrease in nuclear-localized pMad suggests that in addition to its ability to signal as a paracrine factor glial-Gbb appears to act as an autocrine factor to activate BMP signaling in glia.

**Figure 4:**
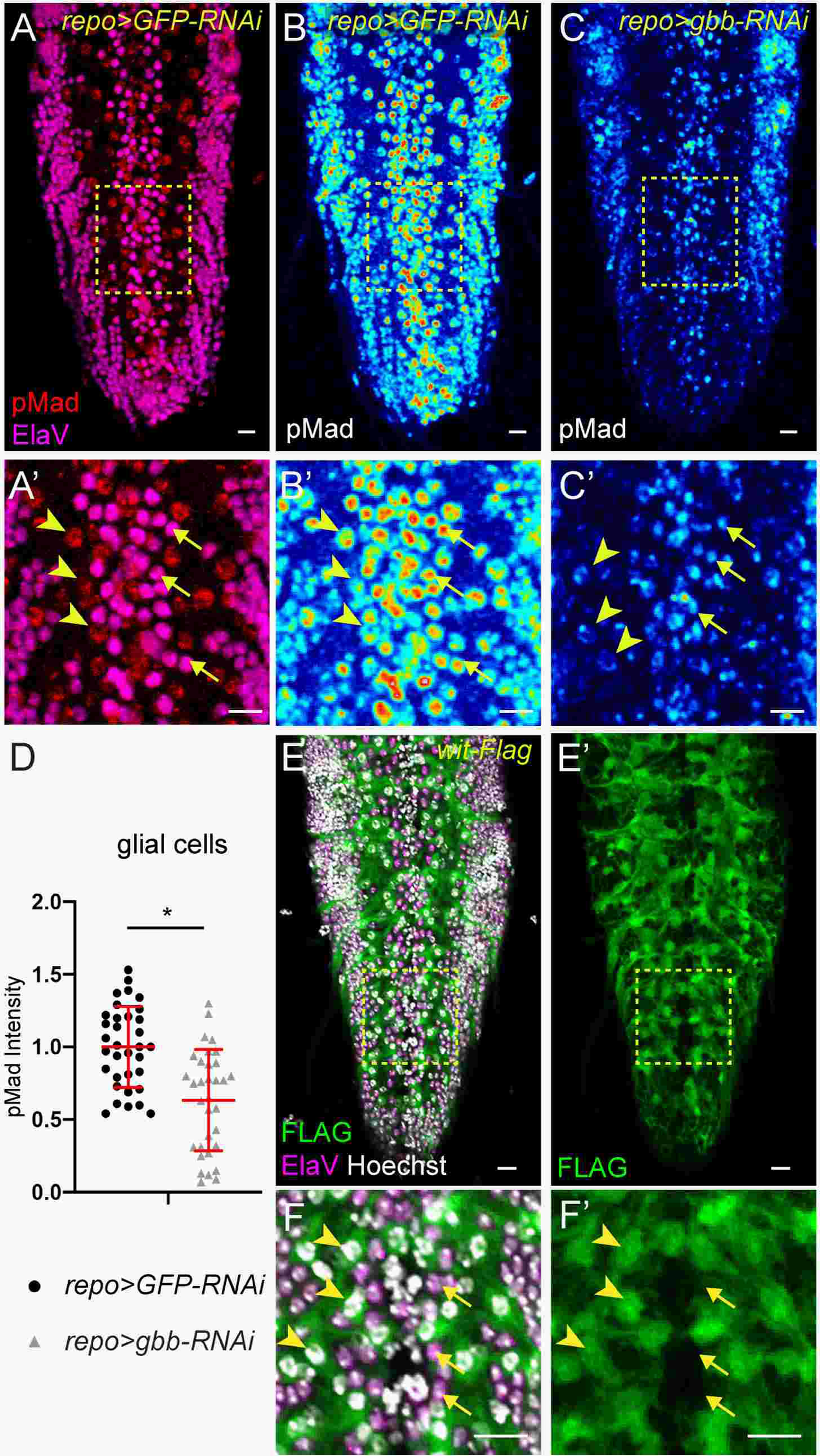
Autocrine activation of BMP signaling by glial-expressed Gbb (A-B’) VNC from a *repo>GFP-RNAi* third instar larva stained with anti-Elav (magenta), and anti-pMad (red) (A). (A’) high magnification of midline containing motor-neurons that innervate segments A5 and A6 located inside the yellow dashed box. Motor neuron nuclei (yellow arrows), glial cell nuclei (yellow arrowhead). (B) VNC and (B’) high magnification of A5 and A6 midline containing motor-neurons from a *repo>GFP-RNAi* third instar larva stained with and anti-pMad (royal). Motor neuron nuclei (yellow arrows), glial cell nuclei (yellow arrowhead). (C) VNC and (C’) high magnification of A5 and A6 midline containing motor-neurons from a *repo>gbb-RNAi* third instar larva stained with and anti-pMad (royal). Motor neuron nuclei (yellow arrows), glial cell nuclei (yellow arrowhead). (D) Quantification of pMad fluorescence intensity in glial cells adjacent (i.e., yellow arrowhead in B’) to A6 motor neuron cluster. Red horizontal bar represents the average and red error bars represent the standard deviation, * shows statistical significance with a T test if p<0.05. (E-E’). VNC from a *Wit-2xTY1-sGFP-V5-BLRP-3x-FLAG* third larval instar stained with anti-Elav (magenta), Hoechst (gray), and anti-Flag (green). (F, F’) are a high magnification of area located inside the yellow dashed box. motor-neuron nuclei (yellow arrows), glial cells nuclei (yellow arrowhead). Scale Bars: 20um

The BMP type II (BMPR2) encoded by *wishful thinking* (*wit*) is required for NMJ size, as *wit* mutants exhibit NMJs with reduced bouton number and overall area (Aberle et al. 2002; Lee et al. 2016; Marques et al. 2002). *wit* mutants lack pMad nuclear localization in neurons of both the embryonic and larval nerve cord (Aberle et al. 2002; Marques et al. 2002), indicating that *wit* function is required for active BMP signaling during these stages. In the embryo (Stage 17), *wit* was shown to be expressed in specific sets of neurons, ie VUM and RP2 (Aberle et al. 2002; Higashi-Kovtun et al. 2010; Marques et al. 2002). In third instar larvae, *in situ* hybridizations show that *wit* is expressed most prominently in the brain lobes (Aberle et al. 2002), with low level diffuse anti-Wit staining in the VNC (Higashi-Kovtun et al. 2010). In order to better understand the expression and localization of Wit receptor in the third instar larval VNC, we made use of a *Drosophila* strain containing a genomic fosmid transgene that encodes a tagged Wit protein (Wit-2xTY1-sGFP-V5-BLRP-3x-FLAG; (Sarov et al. 2016)). We found Wit is localized predominantly to non-neuronal glial cells which do not stain for the pan-neuronally expressed Elav protein (Figure 4E,F’), with comparatively low or little expression of Wit in neurons. *wit* has been shown to be required for BMP signaling activation in the nervous system (Kim and Marques 2012; Marques et al. 2002) and the presence of Wit in glia is consistent with mediation of an autocrine Gbb signal to promote BMP signaling in these non-neuronal cells.

### Expression pattern of *dpp*, the *Drosophila* BMP2/4 ortholog, is distinct from *gbb*

Smad-dependent BMP signaling can be activated by different BMP ligands. To clarify possibility that other sources of BMP ligands exist in the VNC, we examined the expression pattern of the Drosophila BMP2/4 ortholog encoded by *dpp.* The *dpp[blk]-Gal4* transgene expresses GAL4 under the control of disk-region cis-regulatory sequences, reflecting endogenous *dpp* domains of expression (Sarkar et al. 2018; Staehling-Hampton et al. 1994). As indicated by *dpp[blk]-Gal4;UAS-GFP* (*dpp[blk]>GFP*), *dpp* is expressed in a specific set of neurons in the third instar larval VNC (Figure 5). In *dpp[blk]>GFP*, GFP is detected in a distinct set of neurons within the VNC that correspond to the aCC and RP2 motor neurons (Figure 5A). The cell bodies of aCC and RP2 motor neurons occupy invariant positions within a cluster of 12 midline neurons per segment (Doe et al. 1988; Landgraf et al. 1997). The pair of aCC and RP2 motor neurons innervate dorsal internal muscle 1 and muscle 2, respectively, on each side of the larva. The transcription factor Even-skipped (Eve) is expressed in aCC and RP2 motor neurons, in addition to pCC neurons, which do not express *dpp* (Figure S3A (Doe et al. 1988; Garces and Thor 2006)). We further verified the identity of aCC and RP2 motor neurons by expressing membrane bound GFP under the control of the *dpp[blk]-Gal4* driver and tracing their axonal projections to muscles 1 and 2 at NMJ1 and NMJ2, respectively (Figure 5B, E and F).

**Figure 5:**
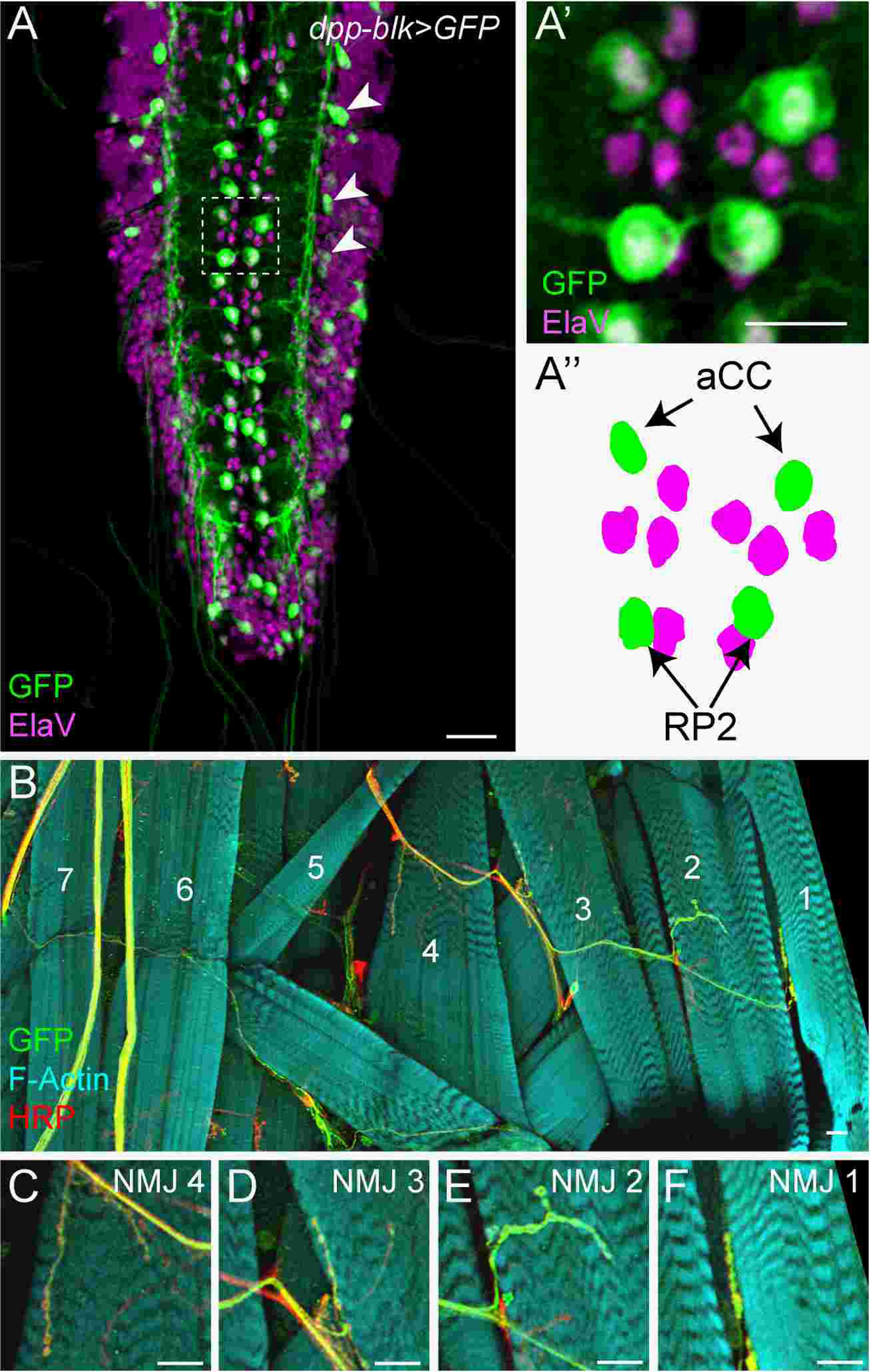
*dpp* is expressed in aCC and RP2 motor-neurons. (A) Third instar larval VNC *dpp-blk-Gal4>UAS-GFP* stained with anti-GFP (green) and anti-ElaV (magenta). Dashed box outlines cluster of midline motor neurons innervating the A3 segment expanded in (A’). (A’’) Position of motor neuron nuclei in A’ image. aCC and RP2 express *dpp* (green), other motor neuron nuclei identified by anti-ElaV (magenta). (B) Internal layer of body wall muscles (numbered) in right A3 hemi-segment of *dpp-blk>GFP* third instar larva stained with phalloidin (cyan). Projections of motor neurons expressing *dpp* marked with GFP (anti-GFP, green). anti-HRP (red). (C-F) Enlarged NMJ 4 (C), NMJ 3 (D), NMJ 2 (E) and NMJ 1 (F). All Scale bars=20um except A’ = 10um

In addition to aCC and RP2, *dpp[blk]>mcd8-GFP* shows a strong signal in 2 to 4 ElaV-positive cell clusters in the lateral VNC (arrowheads, Figure 4A). These clusters correspond to the known location of U motor neurons that innervate the dorso-lateral-internal muscles (Landgraf et al. 1997). Specific markers for U motor-neurons are not available, but in larval filets of *dpp[blk]*>*mcd8-GFP* the projections of these neurons could be traced to their expected point of innervation at NMJ 3 and NMJ4 on dorso-lateral-internal muscle 3 and 4 (Figure 5B-D), indicating that *dpp* is expressed in motor neurons that innervate internal dorsal and dorsal-lateral muscles 1-4.

### Motor neuron-expressed Dpp elicits autocrine BMP signaling required for NMJ growth

Knockdown of *dpp* in its endogenous expression domain in aCC, RP2 and U motor neurons (*dpp[blk]>dpp-RNAi*) results in reduced pMad in nuclei of aCC, RP2 and U motor neurons (Figure 6). pMad fluorescence intensity in aCC (arrowheads) and RP2 (arrows) motor neuron nuclei is 78% of controls (n=5, p=0.03, Figure 6A-B’’, D-E’’, G), and in the U neuron pMad levels are reduced to 68% of the control (n=5, p=4E-03, Figure 6 A-A’’, C-D’’, F-F’’, H). Despite the secreted nature of BMP ligands, activation of BMP signaling based on pMad level is unchanged in motor neuron nuclei adjacent to aCC, RP2 and U when *dpp* is knocked down (n=5, p=0.11, Figure 6I). This finding indicates that while Dpp acts in the autocrine activation of BMP signaling in the aCC, RP2 and U motor neurons, Dpp does not appear to act in neighboring neurons to activate signaling. However, Dpp produced by aCC and RP2 neurons does contribute to pMad-dependent signaling in adjacent glial cells (Figure 6J). The nuclear fluorescence of pMad is decreased to 74% of the control in *dpp[blk]>dpp-RNAi* (p=8.5e-03, n=5). Indicating that Dpp produced in the neurons can also act as a paracrine factor to influence BMP signaling in glia.

**Figure 6:**
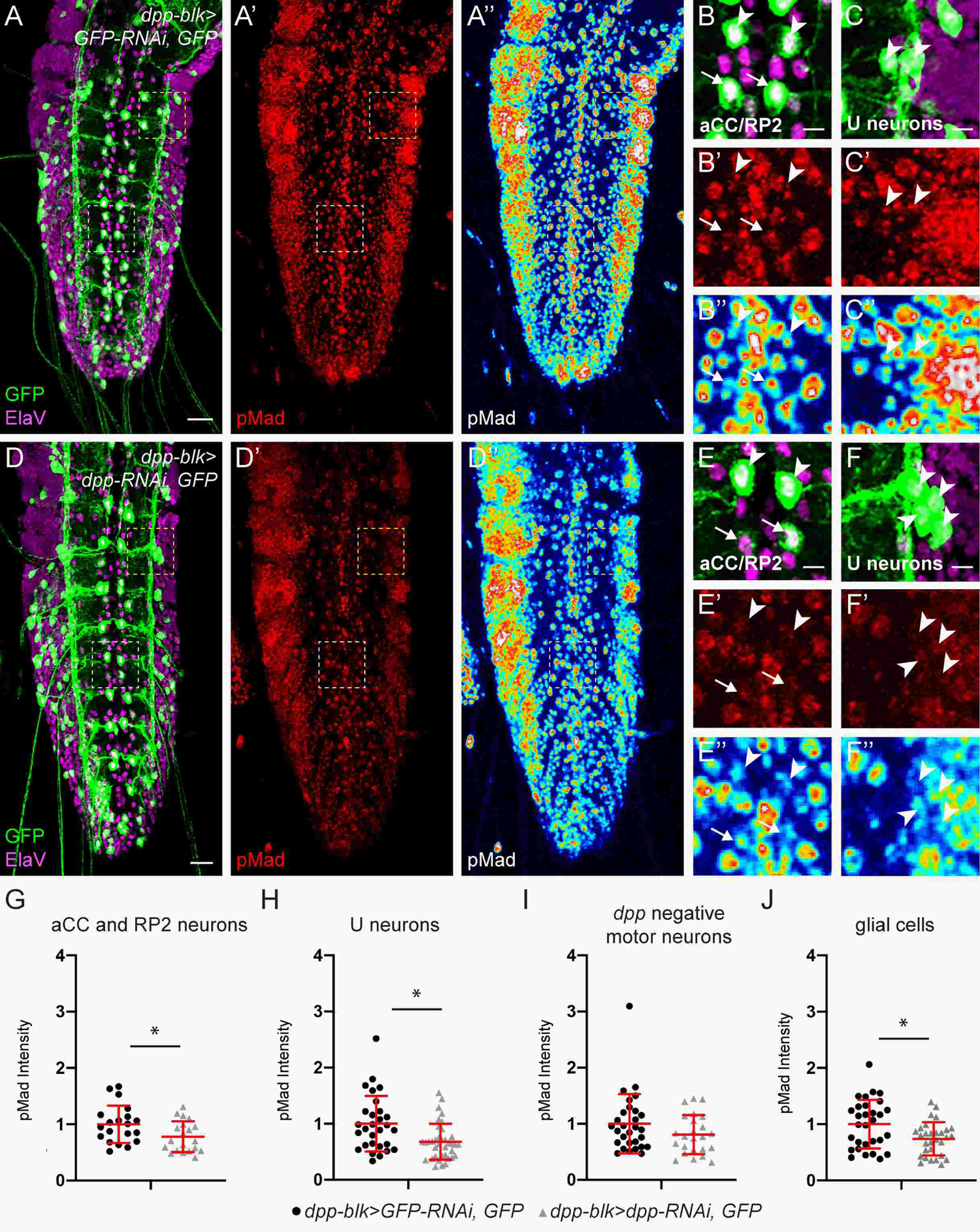
Dpp elicits autocrine signaling in aCC, RP2 and U neurons and paracrine signaling in glial cells. (A-C’’) VNC from a *dpp-blk>GFP* third larval instar stained with (A) anti-GFP (green) and anti-Elav (magenta) and (A’, A’’) anti-pMad. White and yellow dashed boxes mark the midline cluster of motor neurons innervating A5 and the lateral cluster of motor-neurons, respectively, used to quantify pMad fluorescence in the different neuron types, Scale bars=20um. (B-B’’) High magnification of the neurons cluster marked with the dashed white box, Scale bars=5um. (B) anti-GFP (green), anti-ElaV (magenta). (B’, B’’) anti-pMad in red (B’) or with royal lut (B’’). (C-C’’) High magnification of the neurons cluster marked with the dashed yellow box Scale bars=5um. (C) anti-GFP (green), anti-ElaV (magenta). (C’, C’’) anti-pMad in red (C’) or with a royal lut (C’’). (D-D’’) VNC from a *dpp-blk-Gal4;UAS-dpp-RNAi (shdpp2BHB), UAS-GFP* third instar larva, (D) anti-GFP (green), anti-Elav (magenta) and anti-pMad (red D’, royal lut D’’), Scale bars=20um. White and yellow dashed boxes mark the midline cluster of motor neurons innervating A5 and the lateral cluster of motor-neurons used to quantify pMad fluorescence in the different neuron types. (E-E’’) High magnification of neuron cluster marked with white dashed box, Scale bars=5um. (E) anti-GFP (green), anti-ElaV (magenta). (E’, E’’) anti-pMad red (E’) or royal lut (E’’). (F-F’’) High magnification of neuron cluster marked with yellow dashed box, Scale bars=5um. (F) anti-GFP (green), anti-ElaV (magenta). (F’, F’’) anti-pMad red (F’) or royal lut (F’’). (G, H) Quantification of pMad intensity in (G) aCC and RP2 motor, (H) U neuron cluster, (I) midline motor-neurons that do not express *dpp*, and (J) glial cells adjacent to midline motor neurons innervating segments A5 and A6. (G-J) error bars represent standard deviation, * shows p<0.05 based on T test.

Consistent with reduced levels of nuclear pMad in motor neurons of *dpp[blk]>dpp-RNAi*, the number of boutons comprising synapses innervated by RP2 and U neurons, NMJ2 and NMJ3, respectively, are reduced when *dpp* is knocked down in (Figure 7; NMJ2: X⍰= 12.6 boutons in *dpp[blk]>dpp-RNAi* compared to X⍰= 15 in *dpp[blk]>GFP-RNAi* control, n=7, p=0.04; NMJ3: X⍰= 10 boutons when *dpp* is knocked down compared to X⍰=12.4 in *dpp[blk]>GFP-RNAi* control at NMJ3, n=7, p=0.03). Interestingly, this requirement for Dpp-initiated signaling is observed at NMJ2 and NMJ3 despite the presence of Gbb expression in glia and our demonstration that it contributes to NMJ2 and NMJ3 growth (Figure 1 and 3). In an analogous vein, the sole knock down of glial-expressed Gbb is sufficient to compromise the growth of NMJs 2, 3, and 4 (Figure 3A), despite the presence of autocrine signaling initiated by Dpp expressed in aCC, RP2 and U neurons. *dpp* function does not appear essential for NMJ4 growth, as the number of NMJ4 boutons do not change when *dpp* is knocked down (average of 18.2 boutons at NMJ4 in *dpp[blk]>dppRNAi* compared to 18.7 boutons in the *dpp[blk]>GFP-RNAi* control n=7, p=0.79).

**Figure 7:**
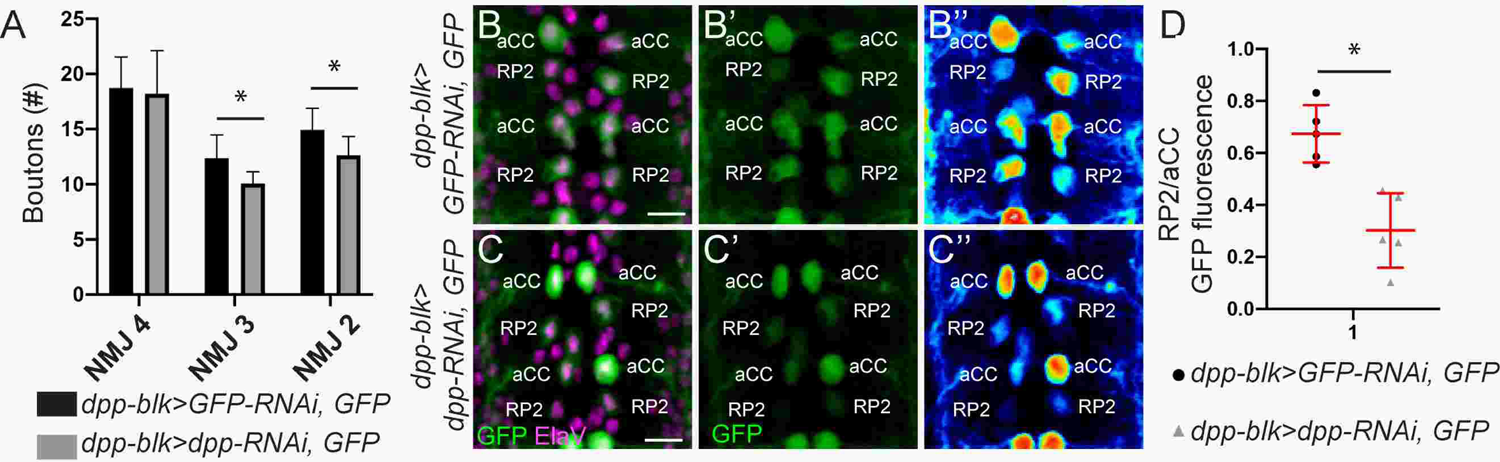
*dpp* is required for growth dorsal NMJ X &Y and autoregulates its expression. (A) Bar chart representing the averaged boutons number at the VNC innervating muscles 4, 3 and 2 of *dpp-blk>GFP-RNAi* controls in black and *dpp-blk>dpp-RNAi* in grey. (B-C’’) Clusters of midline motor neurons innervating A5 and A6 segments of (B-B’’) *dpp-blk-Gal4;UAS-GFP-RNAi,UAS-GFP* and (C-C’’) *dpp-blk-Gal4;UAS-dpp-RNAi,UAS-GFP,* anti-GFP (green), anti-ElaV (magenta). (B’’,C’’) Royal lut shows intensity of GFP signal. Scale Bar = 10uM. (D) Ratio of the GFP signal intensity of RP2 neurons over aCC neurons in the A5 clusters of *dpp-blk>GFP-RNAi* and *dpp-blk>dpp-RNAi* animals. (D) error bars indicate standard deviation, * shows p<0.05 based on T test.

The presence of UAS-GFP in the *dpp[blk]-Gal4* strain reports on the expression of *dpp*. We found that knocking down *dpp* in its own expression domain (*dpp[blk]-Gal4 UAS-dpp-RNAi UAS-GFP*) resulted in a differential response in GFP expression as a reflection of *dpp* transcription, in RP2 versus aCC neurons (RP2:aCC) (Figure 7B-D). *UAS-GFP-RNAi* is used as a control for RNAi-directed gene knock down, and despite driving *GFP-RNAi* in the context of *dpp[blk]-Gal4 UAS-GFP*, sufficient GFP signal remained allowing the identification of cells that normally express *dpp*. The ratio of GFP fluorescence, as a reporter of *dpp* transcription, was determined in control and experimental VNCs and found to be significantly reduced in RP2 motor neurons compared to aCC motor neurons when *dpp* expression is knocked down (Fig 7D). This preferential decrease in GFP signal in RP2 neurons in response to the knock down of *dpp* function, is suggestive of a *dpp*-mediated positive feedback loop that exists in RP2 neurons but not in aCC neurons. An alternative interpretation to the difference in GFP signal could be that RNA interference targeting *GFP* RNA in the control is less effective in aCC neurons, thus skewing the RP2:aCC ratio in controls versus *dpp* knock-down VNCs. To differentiate between these two possibilities, we measured directly the effect of blocking BMP signaling required for the possible autoregulation of *dpp* transcription via pMad, by expressing *Mad-RNAi* (*dpp[blk]>Mad-RNAi*). In this case, *dpp* transcription as reported by *dpp[blk]>GFP*, is dramatically reduced, specifically in RP2 neurons, compared to aCC neurons (Figure S3). The cis-regulatory sequences mediating this autoregulation of *dpp* transcription in RP2 neurons must fall within the disk region of the *dpp* locus (cis-regulatory region contained within *dpp[blk]-Gal4*) (Masucci et al. 1990; Staehling-Hampton et al. 1994). Elucidation of a positive autoregulation of dpp expression in one neuron (RP2) versus another (aCC) highlights the likelihood that the amount of ligand production is tightly regulated between these two cell types, reinforcing the importance of loss-of-function studies to reveal the true function of cell-cell signaling.

## DISCUSSION

### Glia-initiated BMP signaling is important in nervous system development

Glia are well known as critical regulators of the nervous system, contributing to many aspect of the physiological state of neurons and their function (Bittern et al. 2020). Our data show that the BMP, Gbb, is expressed in several types of glia that encase the Drosophila larval VNC and the motor and sensory axonal projections of the segmental nerves, the critical conduit between the CNS and body wall muscles in the control of locomotor activity (Figure 1, S1). Through loss of function studies, we find that glial expressed Gbb contributes to activation of BMP signaling in motor neurons (Figure 8). It is required for attaining normal numbers of boutons in synaptic growth, as well as maintaining larval muscle membrane capacitance, but the knock down of *gbb* in glia did not alone affect spontaneous or evoked neurotransmitter release (Figure 2, 3). This role for synaptic function has been attributed to neuron-expressed gbb. Despite a demonstration that *gbb* knock down in motor and sensory neurons in the context of *gbb* null heterozygotes (*gbb^1^/+; D42-Gal4>UAS-gbb-RNAi*) is associated with presynaptic defects (Hoover et al. 2019; Sanyal 2009), we were unable to detect definitive expression of *gbb* in motor neurons, It is possible that the level of *gbb* expressed in motor neurons is quite low, and given that a *gbb^1^/+* heterozygote would represent a partial loss of *gbb* from all sources, the presynaptic defects observed in *gbb^1^/+; D42>gbb-RNAi* could reflect a partial loss of glial-Gbb combined with a reduction of *gbb* in D42-Gal4/*Toll6*-positive neurons (Sanyal 2009).

**Figure 8:**
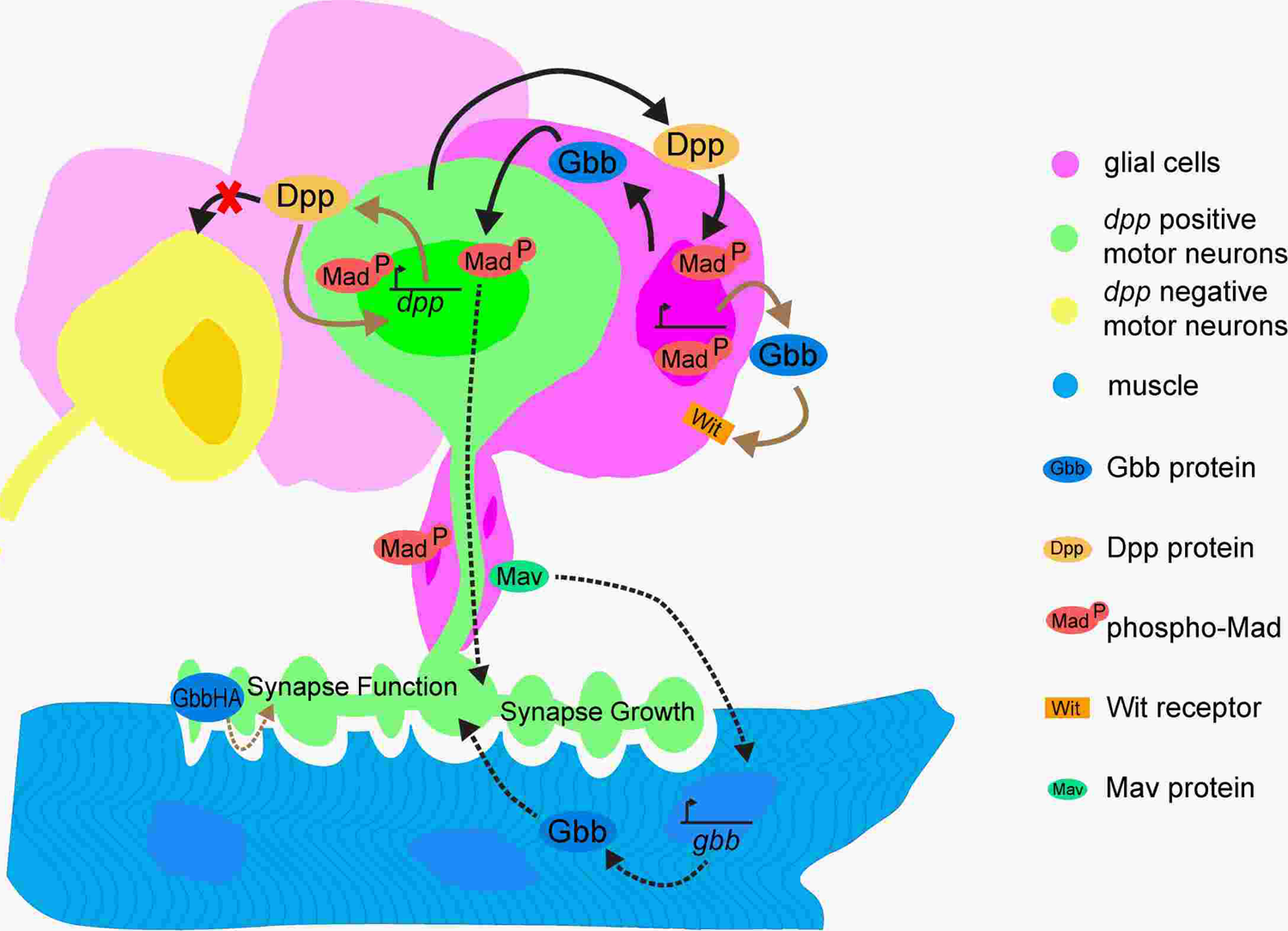
Summary schematic of BMP signaling in Drosophila larval motor circuit. Signaling between glia and neurons initiated by BMP ligands (ovals) demonstrated in this study indicated by solid arrows. Dotted arrows reflect previously published work. Gbb produced by glia (pink) induces pMad-mediated signaling in motor neurons (green) via paracrine signaling (solid black arrow), and in glia via autocrine signaling (solid beige arrow). aCC and RP2 midline motor neurons (green) express Dpp inducing pMad in aCC and RP2 neurons via autocrine signaling (beige line) and glia via paracrine signaling (solid black arrow), but fails to signal (red X) to neighboring motor neurons (yellow). *dpp* transcription in RP2 neurons is positively autoregulated. Presence of pMad in motor neuron nuclei correlates with the bouton number in NMJ (synapse growth). Previous studies have shown glial-expressed Mav regulates expression of *gbb* in muscle (cyan) and when expressed in motor neurons Gbb-HA influences transmission (synaptic function).

In addition to acting in a paracrine manner to activate BMP signaling in both midline motor neurons and peripheral neurons, glial-expressed Gbb also contributes to autocrine activation of BMP signaling in glia (Figure 2, 4, 8). Consistent with this finding, we found as opposed to its earlier neuron-specific expression in the embryonic nervous system, the Wit type II receptor is prominently expressed in glia throughout the larval VNC, with little detectable expression in motor neuron cell bodies (Figure 4). Wit, in glia, likely receives autocrine signals from Gbb, as well as paracrine signals from Dpp from the specific sets of neurons in which it is produced in the larval VNC (Figure 6). Neuron-specific Dpp is also required for autocrine signaling with Mad-mediated autoregulation, in what appears to be specific for *dpp* expression in RP2, but not aCC, motor neurons (Figure 7, S3).

While less is known in the vertebrate motor system with regards to BMP signaling and glia-neuron signaling (reviewed in (Hart and Karimi-Abdolrezaee 2020; Kashima and Hata 2018)), like Gbb, *BMP4* expression has been observed in a subpopulation of glial cells, the Schwann cells, as well as in skeletal muscles, where active BMP signaling is known to impact muscle mass (Chou et al. 2013; Sartori et al. 2013; Winbanks et al. 2013). BMPR2, the Wit ortholog, has been implicated in the regulation of myelination, axon guidance, and changes in motor function associated with abnormalities in receptor trafficking seen in various forms of hereditary spastic paraplegia (Dettman et al. 2018; Tsang et al. 2009; Zhao and Hedera 2013). Certainly, in some of its functions in dendritogenesis and axonal growth, BMPR2 acts as a mediator of Smad-independent non-canonical signaling through the activation of LIMK1 ((Kashima et al. 2016; Lee-Hoeflich et al. 2004; Piccioli and Littleton 2014)). As considerable conservation exists between the *Drosophila* and vertebrate nervous system, as well as the BMP signaling pathway components, it will be of interest to extend our findings of glia-neuron signaling in *Drosophila* to vertebrate systems, including the possibility of a role for BMP-LIMK signaling.

Considering previous reports of the signaling activities of other *Drosophila* BMP/TGF-β family members, i.e., Daw and Mav (Augustin et al. 2017; Fuentes-Medel et al. 2012; Parker et al. 2006), it is clear that ligands in this family each exhibit a specific and distinct expression pattern with the potential for intersecting functions. As we report here, even within the BMP family of ligands, roles for Gbb and Dpp in glia and neurons must be tightly regulated to achieve proper synaptic growth and function (Figure 8). In addition, reports on ligands belonging to the Activin/TGF-β branch indicate their potential for intersection with BMP signaling. Daw, for example, is expressed in glia and required for axon pathfinding specifically in the embryo (Parker et al. 2006) when both *gbb* and *wit* are required for pMad-mediate signaling. But by the larval stage Daw’s role in peripheral glia is reduced compared to that of Mav. From peripheral glia, Mav appears to act through the Punt type II receptor to influence muscle expression of *gbb* (Fuentes-Medel et al. 2012) but the relationship between Mav signals and Gbb signals is not yet clear as a role for *mav* in other glia is not known. Mav does not have a clear mammalian ortholog (Nguyen et al. 2000) but shows most similarity to GDF15, Nodal and TGFβ3 and appears to be mediated by the Activin type I receptor Babo to not only influence p-dSmad2 levels, but also those of the “BMP-specific” pMad ((Gesualdi and Haerry 2007; Hevia and de Celis 2013); T. Knight, pers. comm), Further detailed analyses will be of important to uncover the intersection between BMP and Activin signaling in glia and their role in glia-neuron interactions.

One mechanism by which BMP/TGF-β signaling is regulated may involve the differential outputs exhibited by heterodimer versus homodimer ligands (Akiyama et al. 2012; Bangi and Wharton 2006; Kim et al. 2019; Little and Mullins 2009; Neugebauer et al. 2015). Heterodimeric ligands can only be produced when two different BMP/TGF-β genes are expressed in the same cell. In the context of the larval VNC, *dpp* is expressed in a limited number of neurons compared to the broader expression of gbb detected in glia, trachea and dorsal neurohemal organs. As such signaling elicited by Gbb is most certainly primarily as a homodimeric Gbb;Gbb ligand. and based on the expression pattern observed Dpp:Dpp homodimers would be most prominently produced by aCC and RP2 motor neurons. Interestingly, knock down of *dpp* in its endogenous expression domain did not affect pMad-mediated signaling in neighboring motor neurons that do not express *dpp*, but an autonomous reduction in pMad levels is observed in aCC, RP2 and U neurons (Figure 6). This result indicates that Dpp ligands act close to their site of production. Knocking down Gbb has profound effects on pMad-mediated signaling in cells where it is most prominently expressed, as well as in those where it is not, such as neurons (Figure 2 and 4). This response could reflect the ability of Gbb:Gbb homodimers to act farther from their site of expression, as observed in its role in wing disc patterning (Bangi and Wharton 2006). It could also reflect the extensive network of glial cells in which neuron cell bodies are embedded, providing more access for glial-Gbb to reach neurons. At any rate, the fact that the removal of Gbb;Gbb produced by glia has a profound impact on pMad levels in motor neuron nuclei and on synaptic growth, especially in the motor neuron innervating NMJ4 which also expresses endogenous Dpp, highlights the importance of Gbb-mediated glia to neuron signaling. Furthermore, pMad accumulation in motor neuron nuclei has previously been attributed to retrograde Gbb signals from the muscle (Berke et al. 2013; Fuentes-Medel et al. 2012; McCabe et al. 2003). More recently, it has been argued that BMP signaling in the motor neurons of the third instar larval VNC is primarily due to autocrine Gbb signaling, based on the ability of neuron-driven *gbb* (*gbb^1^/gbb^2^; D42-Gal4>UAS-gbb*) but not muscle-driven Gbb expression to rescue the loss of pMad in gbb null larvae (*gbb^1^/gbb^2^*; *BG57-Gal4>UAS-gbb*) (Hoover et al. 2019). In neither genotype is *gbb* expression in glia, lost in a *gbb* null animal (*gbb^1^/gbb^2^*), restored by the overexpression of *gbb* in neurons or muscles. Loss of *gbb* from glia is sufficient to compromise nuclear pMad accumulation in neurons (Figure 2) and while expressing *gbb* in neurons is capable of eliciting pMad-mediated signaling, under endogenous conditions glial-Gbb signals are required to induce BMP signaling in neurons and for proper synaptic growth.

Neuron-glia interactions in a Drosophila Parkinson’s model have implicated the activation of BMP signaling in glia of the larval brain in the age-dependent neurodegeneration of DA neurons associated with LRRK2 overexpression (Maksoud et al. 2019). Similarly, in a paraquat-induced model of Parkinson’s disease, the knockdown of Mad in glia was found to protect against the toxic effects of paraquat on DA neuron survival and adult longevity (Maksoud et al. 2019). The different response of neurons and glia to Gbb expression in the brain versus the VNC could depend on a number of factors including cell-specific expression of different receptor complex components, i.e., different type I and type II receptor types, and/or differential processing of the ligand, as found to be the case in CCAP neurons (Anderson and Wharton 2017). Furin 1, a pro-convertase important in modulating LRRK2’s ability to regulate synaptic transmission (Penney et al. 2016), proteolytically processes BMP proproteins to generate mature ligands. The ability of Fur1 to cleave at specific sites in Gbb is influenced by O-glycosylation and thus, different ligand forms may be secreted from cells depending on glycosylation (Anderson and Wharton 2017). It is intriguing that Fur1 has been identified as a component whose function modulates the pathogenic outcome of a LRRK2-associated Parkinson’s model. The predominant form of Gbb produced by glia, brain neurons and muscles is not known and it may differ, as in CCAP neurons, where the active Gbb ligand form was shown to exhibit differential outcomes and receptor binding preference (Anderson and Wharton 2017).

Gbb exhibits multiple functions during nervous system development and homeostasis. To some extent its functions in synaptic growth and function are separable and thought to reflect different sources of Gbb. The differential activation of BMP signaling is also reflected by multiple pools of pMad. In addition to pMad accumulation in the nuclei of muscles, some neurons and glia, pMad is also evident at the NMJ (Dudu et al. 2006). Synaptic pMad is associated with clustering of glutamate receptors (GluRs) and synaptic stability but not synaptic growth (Sulkowski et al. 2014). Consistent with roles for *gbb*, synaptic pMad does not depend on *gbb* (Sulkowski et al. 2016), as we find that endogenously expressed glial-Gbb is required for synapse growth but alone its downregulation does not affect synapse function. However, it has been shown that when expressed in neurons, Gbb-HA secreted into the synaptic cleft, associates with the α_2_δ-3 auxiliary calcium channel sub-unit which acts as an extracellular scaffolding factor to maintain synapse function (Hoover et al. 2019). It is possible that Gbb’s role in regulating neurotransmitter release may depend solely on Gbb released into the synaptic cleft, or glial-Gbb could also contribute.

### The expression of Dpp is under the control of a positive feedback loop

The necessity for tight regulation of BMP signaling is evident as the underlying cause of a number of human developmental abnormalities. Indeed, over-activation or a failure in the maintenance of signaling is responsible the induction of heterotopic bone formation in fibrodysplasia ossificans progressiva (FOP), disorders of craniofacial development, and the loss of germline stem cell populations (Graf et al. 2016; Kaplan et al. 2009; Shore et al. 2006; Wagner 2007). Regulation of the pathway may begin with the spatial and temporal regulation of ligand gene expression as a means to ensure continued supply of a ligand. Our results indicate that *dpp* expression is under the control of positive autoregulation by cis-regulatory sequences located in the disk region the *dpp* locus. We find that *dpp[blk]*-reported expression of *dpp* transcription in RP2 neurons is compromised in a Mad-dependent manner when *dpp* is knocked down by RNAi. Dpp’s ability to positively autoregulate its own expression has also been seen in the embryonic visceral mesoderm and with specific reporter constructs in the developing wing disc (Galeone et al. 2017; Hepker et al. 1999; Staehling-Hampton and Hoffmann 1994). Similarly, in immortalized mouse osteoblasts, a proximal regulatory sequence located in the 5’ of the *BMP2* locus, a *dpp* ortholog, has been shown to be activated by BMP2 (Ghosh-Choudhury et al. 2001). Beyond direct transcription regulation, numerous cases of pathway regulation through the activation of agonists and antagonists that act in the extracellular or cytoplasmic space have been documented for the BMP/TGF-β pathway as mechanisms to maintain or downregulate signaling (Faherty et al. 2016; Huang et al. 2015; Szuperak et al. 2011; Tremml and Bienz 1992; Yu et al. 1996).

In the case of the Drosophila nervous system, we have revealed a role for BMP signaling in glia-neuron communication in synaptic growth. Future studies will aim to understand how this system influences and fully integrates with other signaling requirements and cellular responses required for proper synaptic structure and function.

## ACKNOWLEDGEMENTS

We would like to acknowledge the Bloomington Drosophila Stock Center (NIH P40OD018537), the Vienna Drosophila Resources Center (VDRC, www.vdrc.at), the Developmental Studies Hybridoma Bank (DSHB), created by the NICHD of the NIH and maintained by The University of Iowa, Department of Biology, for providing critical strains for our analysis, as well as the wealth of information provided by Flybase (supported by NIH-NHGRI U41 HG000739, the British Medical Research Council MR/N030117/1). This work was supported in part by funds to K.A.W. from NIH GM068118, The Judith and Jean Pape Adams Foundation, and the ALS Finding a Cure Foundation. T.K. received assistance from the Brown University Undergraduate Research & Teaching Award and A.H. received support from Brown Institute for Brain Science Robin Chemers Neustein Graduate Fellowship.

## Supplemental Figures

**Figure S1:** (A-A’) VNC of a *repo-Gal4;UAS-mcd8-GFP* (*repo>mcd8GFP*) and (B-D) *gbb-Gal4;UAS-mcd8-GFP* (*gbb>mcd8GFP*) third instar larvae stained with anti-GFP (green) and anti-Repo (magenta). Dashed line indicates position of cross-section in A’ and B’. (A,A’) Extensive network of glia encasing the VNC and axonal projections of motor and sensory neurons in the segment nerves is evident. (B) *gbb* is expressed in muscles (asterisk), dorsal neurohemal organs (white arrowhead), glia encasing the VNC as shown in cross section (B’, yellow arrowhead marks edge of trachea), peripheral glia in segmental nerves, and trachea (thinner strands, also shown in C). *gbb-Gal4-#5* is an independent transgene insertion than that shown in Figure 1. (C) Posterior extremity of a *gbb#5>mcd8GFP* third larval instar. *gbb* is expressed in glial cells wrapping the axonal projections of the segmental nerves marked by Repo-positive nuclei (white arrows), tracheal cell nucleus lacking Repo staining (yellow arrowhead). (D) *gbb* expression in muscles. Hoechst-stained DNA marks nuclei (white), anti-GFP (green). *gbb-Gal4* strain in this panel is same as in Figure 1. Scale bar = 20um

**Figure S2:** B**M**P **signaling activated in peripheral glia** (A-A”) segmental nerves projections and posterior terminus of VNC from repo-Gal4 third instar larva. (A) anti-Repo, (A’) anti-pMad (PS3), (A”) merged channels. Scale bar = 20um

**Figure S3:** d**p**p **transcription in RP2 neurons is under positive autoregulation** (A-A’’) Two clusters of dorsal midline motor neurons in a *dpp-blk-Gal4;UAS-shdpp2BHB, UAS-GFP* third instar larval VNC stained with anti-GFP (green) and anti-Eve (magenta). aCC, pCC, and RP2 motor-neurons labelled. (B-C’’) Midline clusters of dorsal midline motor neurons innervating A5 and A6 segments of *dpp-blk-Gal4;UAS-GFP-RNAi, UAS-GFP* (B-B’’) and *dpp-blk-Gal4;UAS-Mad-RNAi,UAS-GFP* (C-C’’) stained with anti-GFP (green) and anti-ElaV (magenta). A royal lut shows the intensity of the GFP signal (B’’ and C’’). Asterisks mark RP2 motor-neurons. Scale bar = 5um.

